# Mucosal Immunization of Cynomolgus Macaques with Adenoviral Vector Vaccine Elicits Neutralizing Nasal and Serum Antibody to Several SARS-CoV-2 Variants

**DOI:** 10.1101/2022.02.21.481345

**Authors:** Becca A. Flitter, Colin A. Lester, Sarah N. Tedjakusuma, Emery G. Dora, Nadine Peinovich, Mario Cortese, Clarissa I. Martinez, Clara B. Jegede, Elena D. Neuhaus, Sean N. Tucker

## Abstract

The emergence of SARS-CoV-2 variants continues to be a major obstacle for controlling the global pandemic. Despite the currently authorized SARS-CoV-2 vaccines ability to reduce severe disease and hospitalization, new immunization strategies are needed that enhance mucosal immune responses, inhibit community transmission, and provide protection against emerging variants. We have developed a mucosally delivered, non-replicating recombinant adenovirus vector (rAd5) vaccine, that has proven efficacy in the clinic against other respiratory viruses [1]. Here we evaluated the immunogenicity of three candidate SARS-CoV-2 vaccines in cynomolgus macaques that contained spike (S) and/or nucleocapsid (N) from either the Wuhan or the beta variant to select a candidate for future clinical development. Mucosal immunization with the Wuhan specific S vaccine (ED90) induced significant cross-reactive serum IgG responses against to Wuhan, beta, gamma and delta lineages, and generated substantial serum neutralizing activity. In nasal samples, ED90 immunization induced 1000-fold increases in IgA to all variants of concern tested and had neutralizing activity against Wuhan and delta. While immunization with the beta specific vaccine (ED94) enhanced IgG and IgA responses to homologous beta variant S and RBD, this approach resulted in less cross-reactive responses to other variants in the serum and nasal passages compared to ED90. As ED90 immunization induced the most robust cross-reactive systemic and mucosal antibody responses, this candidate was chosen for future clinical development.

## Introduction

Several SARS-CoV-2 vaccines, all administered by injection, have been authorized and approved. These vaccines all produce substantial serum antibody responses that are potent against the original parental strain. However, antibody titers wane and as new variants of concern (VOC) appear, vaccine induced protection against SARS-CoV-2 infection drops. The AstraZeneca vaccine ChAdOx1 was found to be less protective against mild to moderate disease caused by the beta variant (B.1.351) with an efficacy of only 10% found in one study [2]. The BioNTech/Pfizer mRNA vaccine was estimated to be only 56% effective at preventing delta variant (B.1.617.2) infection after two doses of vaccine, with efficacy improving to 92% only after three doses [3]. Furthermore, two doses of currently approved mRNA vaccines offer limited protection against infection with the omicron variant (B1.1.529) with a third dose providing only 37% efficacy [4]. Studies indicate the omicron variant appears to preferentially replicate in the upper respiratory tract and has substantial mutations in the receptor binding domain [5]. Which suggests that selective pressure by the systemic immune response, ie serum IgG, may have contributed to the emergence of the omicron variant. Vaccine manufacturers are currently beginning trials on variant-specific COVID-19 vaccines. However, given the speed of variant appearances, the continuous production and manufacturing of variant-matched vaccines is unlikely to control the pandemic.

An alternative approach is to create a cross-reactive IgA protective response at mucosal sites where viral entry and primary infection occurs. In mucosal tissues, plasmablasts predominantly produce secretory IgA (SIgA) and due to increased valency, this immunoglobulin has been shown to have better neutralizing activity against invading pathogens compared to IgG [6, 7]. As SIgA can be multimeric, the immunoglobin can directly bind multiple virus particles blocking receptor binding and subsequent entry into target cells (Reviewed by [8]). SIgA also prevents respiratory and enteric pathogens from infecting mucosal sites by immune exclusion, steric hindrance, and by inhibiting intracellular viral production [9-11]. Creating a potent SIgA response to SARS-CoV-2 through mucosal vaccination may greatly reduce both viral shedding and transmission to other individuals [12] and could have a significant impact on the trajectory of the pandemic. The potency of mucosal immunization in blocking aerosol transmission has been demonstrated in a hamster SARS-CoV-2 infection transmission model [13] where unvaccinated hamsters were protected from disease when co-housed with hamsters administered mucosal immunization followed by viral challenge [13]. Therefore, enhancing the production of SIgA through mucosal vaccination could greatly reduce SARS-CoV-2 transmission and is a potential effective future strategy for reducing the number of infections.

We have developed a mucosal vaccine platform that uses a replication-defective human adenovirus type-5 vectored vaccine (rAd5) which expresses a protein antigen along with a novel toll-like receptor 3 (TLR3) molecular agonist adjuvant. The vaccine is formulated to be dried and tableted to and delivered by simply swallowing with a glass of water. Over 500 human subjects have been administered these oral vaccines which are well tolerated, and generate robust humoral and cellular immune responses to the expressed antigens [14-16]. We have previously published data on the immunogenicity and selection of a COVID-19 clinical candidates in hamsters and mice [13, 17], but interpreting results from small rodent models do not always translate in human clinical trials. The original vaccine candidate to be test in phase I clinical trials induced substantial T cell responses in humans, elicited some IgA in the mucosa, but produced no serum neutralizing antibody responses after a single dose.

To better understand the humoral immunological responses to our mucosal SARS-CoV-2 vaccines, we examined three candidates in cynomolgus macaques. The goal was to select a vaccine candidate that could elicit protective mucosal and systemic immune responses to a diverse group of variants, so multiple candidates were evaluated. Our original clinical candidate contained both the parental spike (S) and nucleocapsid (N), and since N may influence immunogenicity to S, we evaluated both S only and S plus N Wuhan vaccine candidates. We also tested a variant specific vaccine approach against the beta, to determine if immunizing with homologous S proteins would provide more protective efficacy when new emerging variants arise. Therefore, we evaluated candidates expressing S only (ED90) and S + N (ED88), and S-only beta-variant specific candidate (ED94). As protein-based vaccines are also being deployed around the world, a protein subunit vaccine prime followed by mucosal adenoviral immunization boost was used a comparator group. We evaluated these vaccine candidates (and one heterologous immunization regiment) to induce immune responses to different variant S proteins (Wuhan, alpha, beta, gamma and delta). As delivering human specific tablets in non-human primates is challenging and often ineffective, we used intranasal vaccination of a liquid formulation in our immunogenicity evaluation. We showed that intranasal vaccine administration elicits both binding and neutralizing IgG in serum similar to what has been reported with other rAd and mRNA vaccines [18, 19]. The most potent vaccine candidate (ED90) tested, generated substantial cross-reactive antibodies to the delta and beta variant S proteins and produced more serum antibodies than other candidates evaluated. Moreover, vaccination with ED90 induced substantial IgA antibody responses in the nasal mucosal of non-human primates. Here we demonstrate that mucosal vaccination with ED90 elicits both serum and mucosal responses to major VOC.

## Results

### rAd5 mucosal vaccination with ED90 elicits strong cross-reactive serum IgG and IgA

To investigate humoral immunogenicity of three clinical vaccine candidates, five groups of cynomolgus macaques were vaccinated with rAd5 containing transgenes of S protein from either Wuhan (ED90) or beta (ED94) lineages, or a combination of Wuhan S and N proteins (ED88) **(Figure 1A)**. Four macaques in each group were immunized intranasally with a prime boost regimen on day 0 and 28 and nasal swab samples and serum were collected at D-1 and D15, D30, D44 and D58 post vaccination **(Figure 1B)**. One group received intramuscular injection prime with recombinant S protein followed by intranasal boost immunization with ED88 **(Figure 1C**). No adverse clinical events were recorded after vaccination and overall body weights increased over the 8-week study **(Sup Figure 1)**.

**Figure 1:**
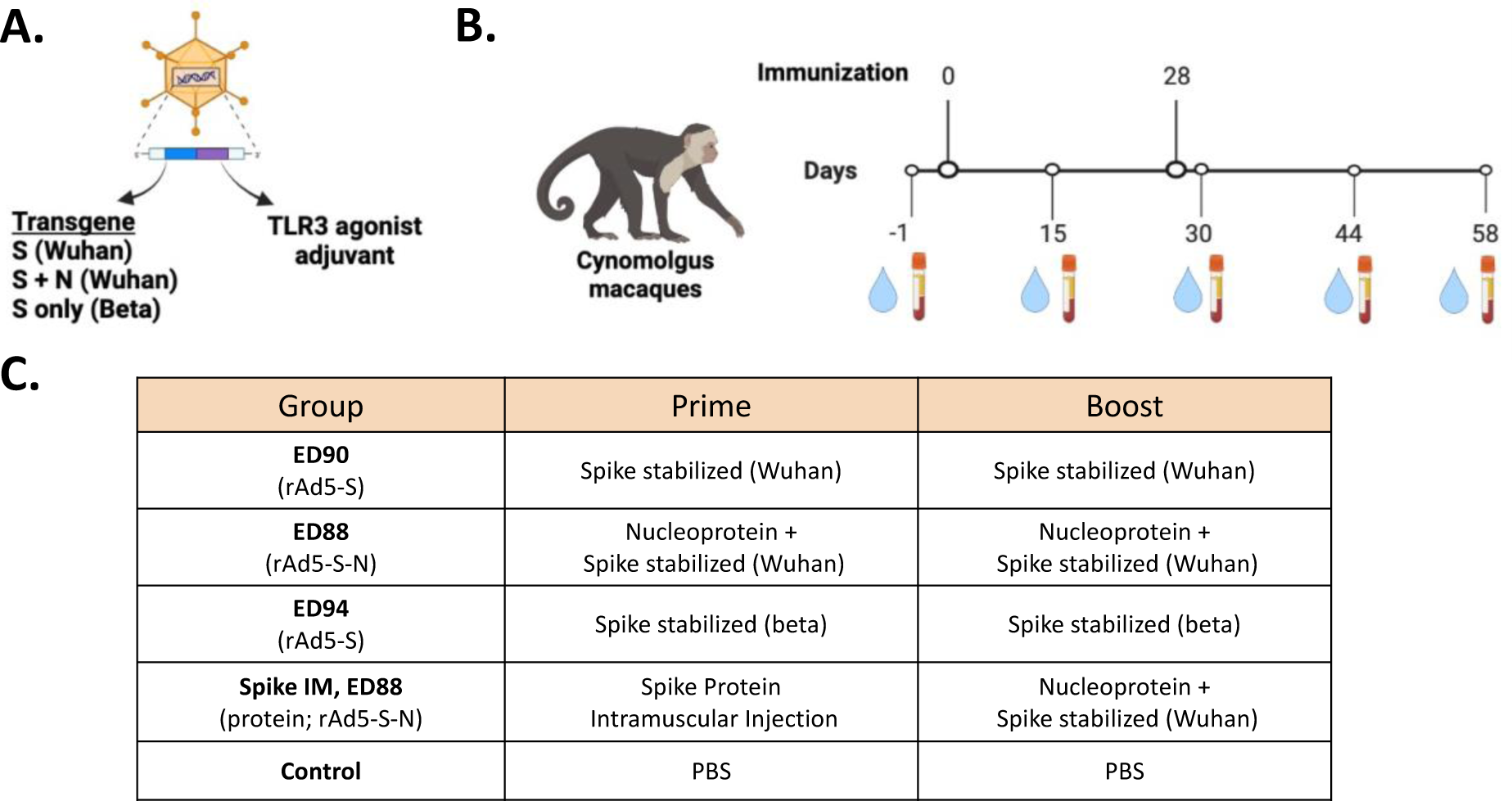
Immunogenicity study design for assessing of mucosal delivery of potential rAd5 vaccine candidates. **(A)** Illustration of rAd5 vector candidates. **(B)** Study timeline for prime boost vaccine administration and serum and nasal swab sample collection. (C) Vaccine candidates evaluated for immunogenicity in cynomolgus macaques.

Two weeks following single vaccine dose, ED90 immunized animals had 2.5 to 3-log increases in serum IgG specific to full length S and RBD proteins from Wuhan and five variant lineages, beta, delta, alpha and gamma **(Figure 2A-F; Supplemental 2)**. Animals receiving ED94 had similar increases of beta and gamma specific IgG to animals vaccinated with ED90, but lower responses to the Wuhan, delta and alpha variants. Therefore, the variant specific vaccine (ED94) was effective at generating IgG responses against homologous or a closely related variant but elicited less cross protective serum IgG to other S specific strains. Animals administered ED88 had moderate increases in anti-S protein IgG responses, compared to ED90 vaccinated animals, which was unexpected at both vaccines contain Wuhan S transgene. Animals primed with purified S protein on D0 followed by intranasal boost at D28 with ED88, had lower serum IgG responses to all RBD and S protein variants. Serum IgA specific to Wuhan, alpha, delta and gamma was generated in animals vaccinated with ED90 **(Figure 3A-F, Supplemental 2)**; which as similar to what was observed with IgG, but slightly lower in magnitude. However, the full length and RBD beta variant specific IgA serum responses were the highest in animals vaccinated with ED94. Interestingly it was observed that some vaccine groups had an increase in serum IgA detected on D44, two weeks after boost, whereas serum IgG levels remained constant. Antibody responses to N were not detected at any timepoint including animals vaccinated with ED88 (data not shown). Overall, these results demonstrate NHPs immunized with vaccine candidate ED90 generate both cross-reactive serum IgG and IgA responses to multiple VOC.

**Figure 2:**
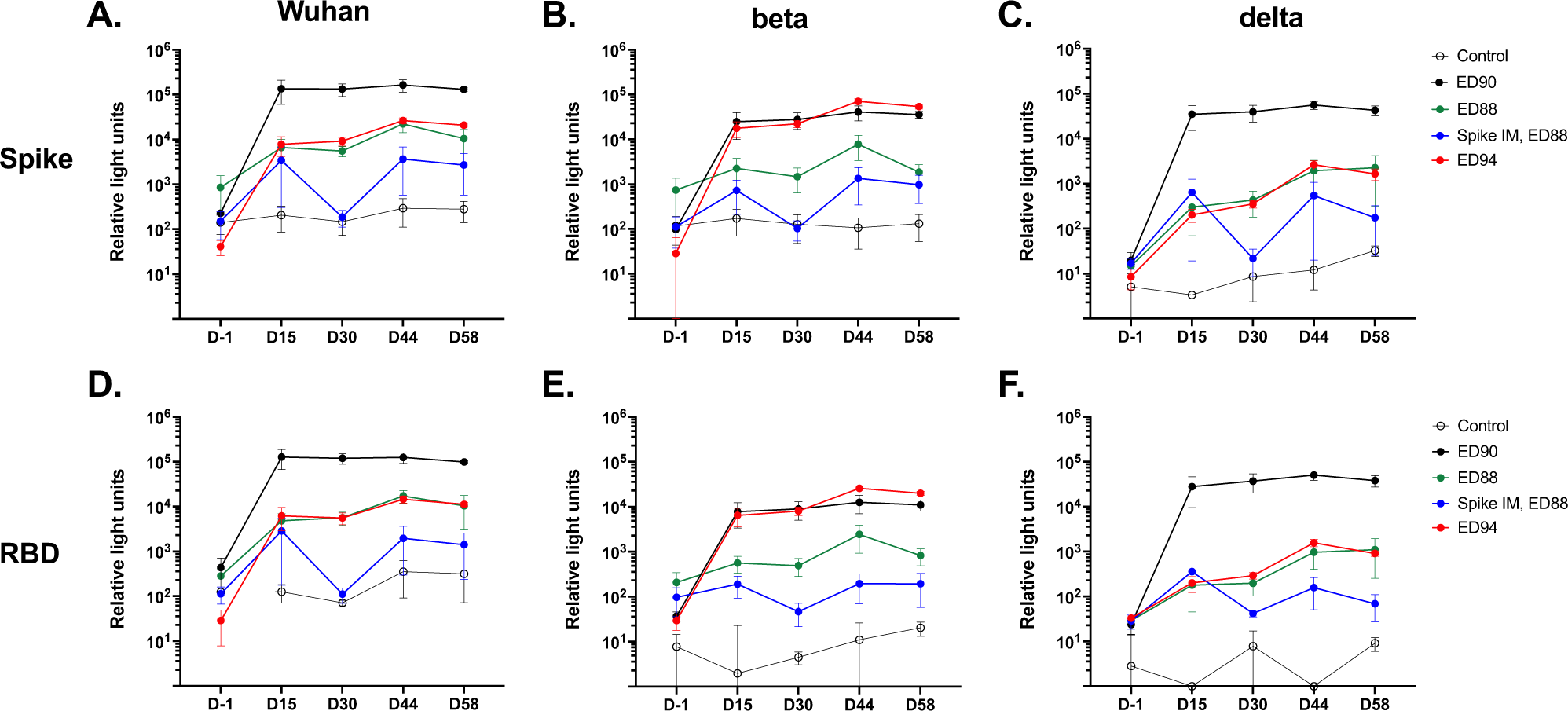
Mucosal immunization with ED90 elicits strong cross-reactive IgG in the serum. Animals were immunized on D0 and D28 with vehicle control (open circles) ED88 (green), ED90 (black), ED94 (red), or primed on D0 with IM delivery of spike protein followed with ED88 boost (blue) on D28. S protein specific serum IgG was measured against **(A)** Wuhan **(B)** beta variant (B.1.351) and **(C)** delta variant (B.1.617.2) on D-1, D15, D30, D44 and D58 using MesoScale Diagnostics (MSD) electrochemiluminescence. Serum IgG against **(D)** Wuhan **(E)** beta and **(F)** delta RBD were measured concurrently. Data expressed as MSD relative light units, SEM (n = 4).

**Figure 3:**
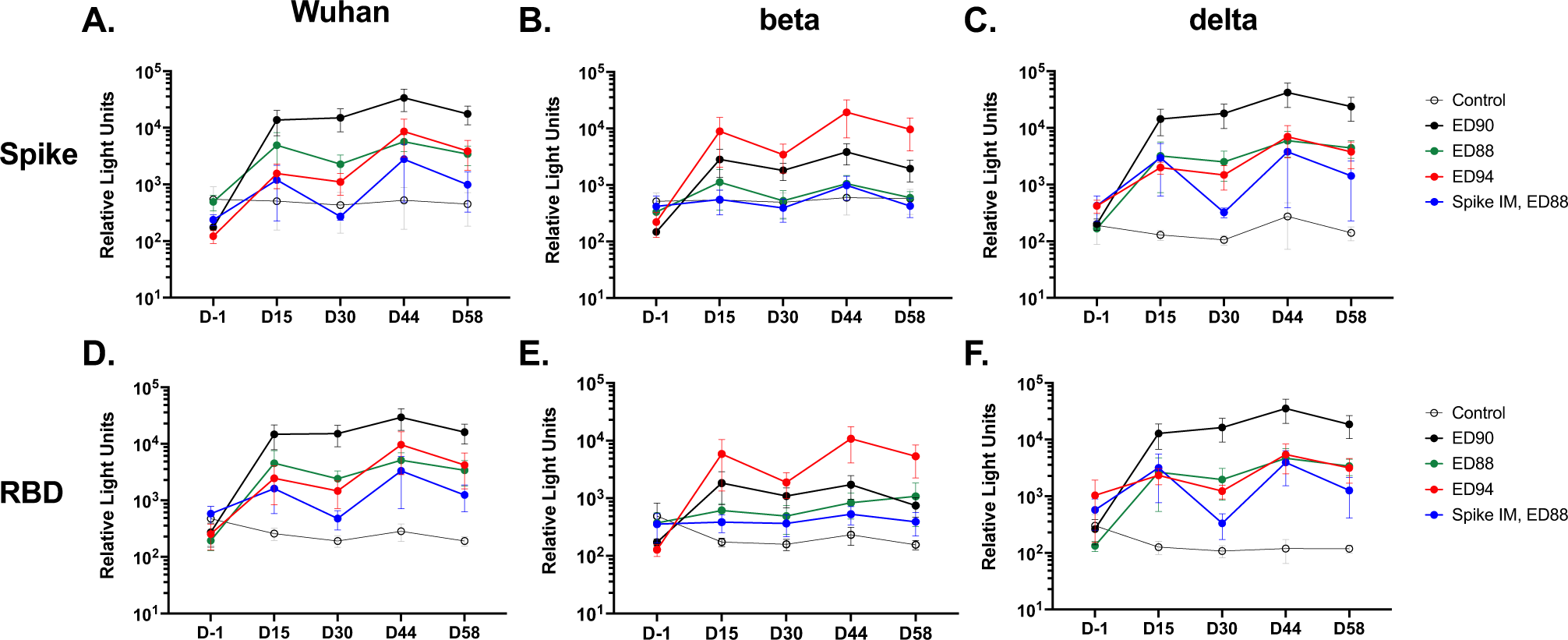
Mucosal vaccination with ED90 elicits strong cross-reactive serum IgA responses. S protein specific serum IgA was measured against **(A)** Wuhan **(B)** beta variant (B.1.351) and **(C)** delta variant (B.1.617.2) on D-1, D15, D30, D44 and D58 using MesoScale Diagnostics (MSD) electrochemiluminescence. Serum IgA against **(D)** Wuhan **(E)** beta and **(F)** delta RBD were measured in parallel. Data expressed as MSD relative light units, SEM (n = 4).

### Immunization with ED90 produces neutralizing serum antibody to Wuhan, beta and delta

To assess antibody functional activity plaque reduction neutralization tests (PRNT) were conducted and IC_50_ titers were determined against SARS-CoV-2 virus strains, WA/2020 (Wuhan), RSA (beta) and delta on NHP serum collected at D-1, D30 and D58 **(Figure 4A)**. By D30, three out of four animals immunized with ED90 had neutralizing titers to the Wuhan virus ≥IC_50_ titer of 50 denoted by the dotted line, the proposed protective titer in a NHP model [20]. By D58 all 4 monkeys vaccinated with ED90 had ≥IC_50_ titer of 50. None of the other vaccine candidates elicited serum neutralizing titers to the Wuhan Strain. All four ED94 vaccinated animals generated neutralizing antibody titer by D58 to the homologous RSA virus. 50% of the ED90 immunized animals had neutralizing activity against the RSA strain by D30, however there was not an increase in titer following boost on D58. Surprisingly, animals immunized with ED90 were the only group to generate neutralizing antibody titers by day D30 against the delta variant. 100% of ED90 vaccinated animals continued to have high protective neutralizing serum antibody on D58. Only 50% of animals immunized with ED88 generated a neutralizing antibody by D58.

**Figure 4:**
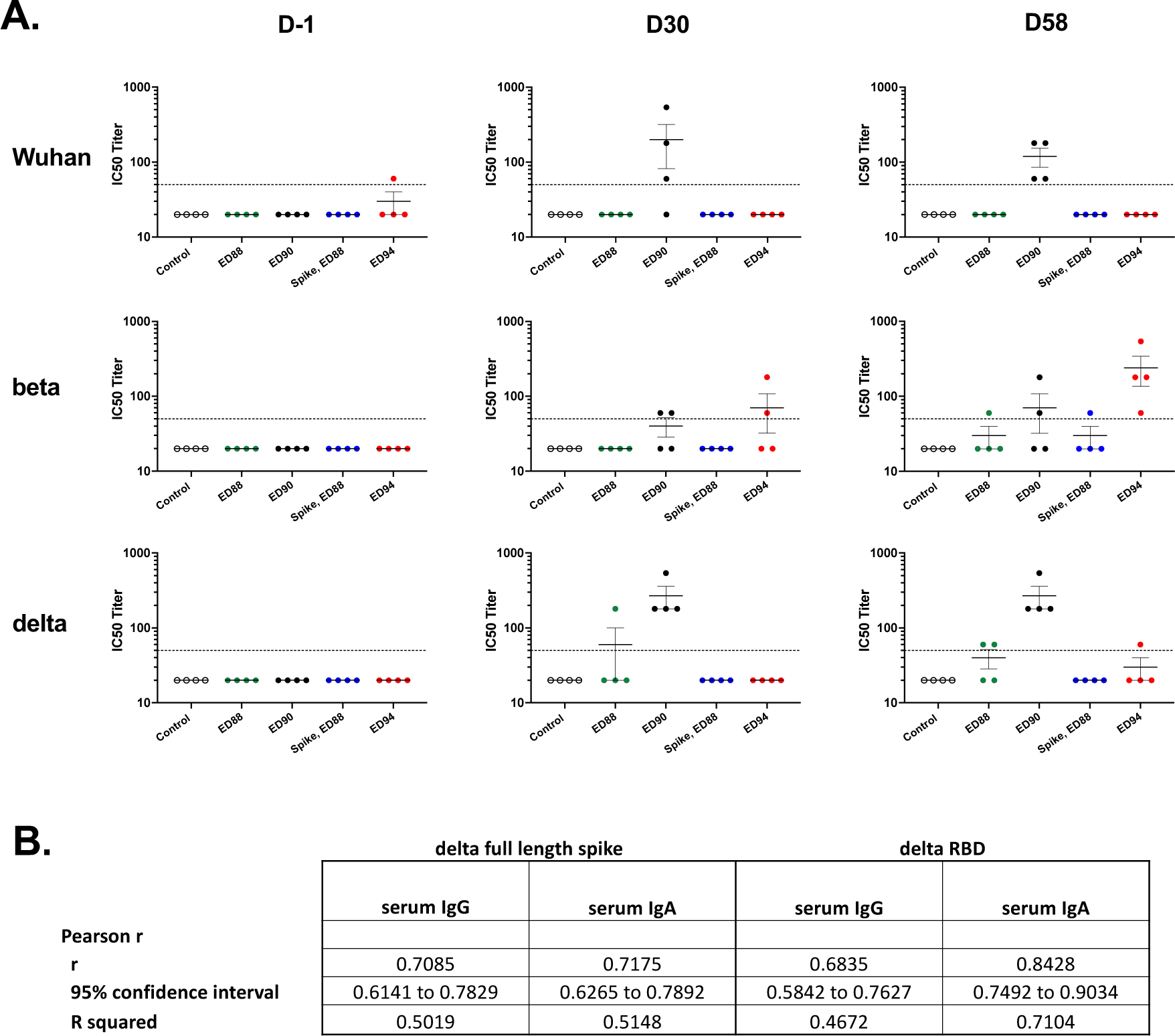
ED90 elicits serum neutralizing IC50 titers to both Wuhan and delta. (A) Serum neutralizing activity against Wuhan, beta or delta SARS-CoV-2 virus at D-1, D30 and D58 by PRNT. Dotted line denotes proposed NHP IC50 protective serum titer (IC50 = 50) Plaque forming units were quantified and IC50 titers reported, SEM (n = 4). (B) Pearson correlation analysis comparing delta variant serum IC50 neutralization titers to total serum IgG and IgA levels.

Surprisingly, animals vaccinated with ED90 had the highest serum IgG and IgA and neutralization activity to the delta strain. By Pearson’s correlation analysis we evaluated if serum IgG and IgA specific to delta were associated with the homologous strain neutralization titers. As expected, both serum IgA and IgG specific to delta RBD correlated with delta PRNT titers, (Pearson coefficient, 0.843 and 0.684 respectively), however, IgA was more highly associated with serum neutralizing activity than IgG **(Figure 4B)**. Similar results were found with Wuhan in ED90 vaccinated animals, confirming serum IgA quantities against the RBD portion correlated more with neutralizing activity in the serum compared to IgG (data not shown).

### The vaccine candidate ED90 elicits cross-reactive nasal IgA specific to S protein and RBD to variants of concern

The production of SIgA at mucosal surfaces has several protective immune functions including contributing to the first line of defense for restricting pathogen entry. We collected secretions by swabbing both nostrils, on D-1, D15, 30, 44 and 58 and measured IgA from the eluted and pooled samples. Nasal IgA responses were reported normalized to total IgA quantified in each respective sample and expressed as fold change. Nasal IgA specific to Wuhan, delta, alpha and gamma full length and RBD proteins all increased nearly 1000-fold, by day 58 in ED90 immunized animals **(Figure 5; Supplemental 3)**. ED94 vaccine candidate elicited the most nasal IgA specific to beta. However, ED90 was able to induce a 200-500-fold increase by D58.

**Figure 5:**
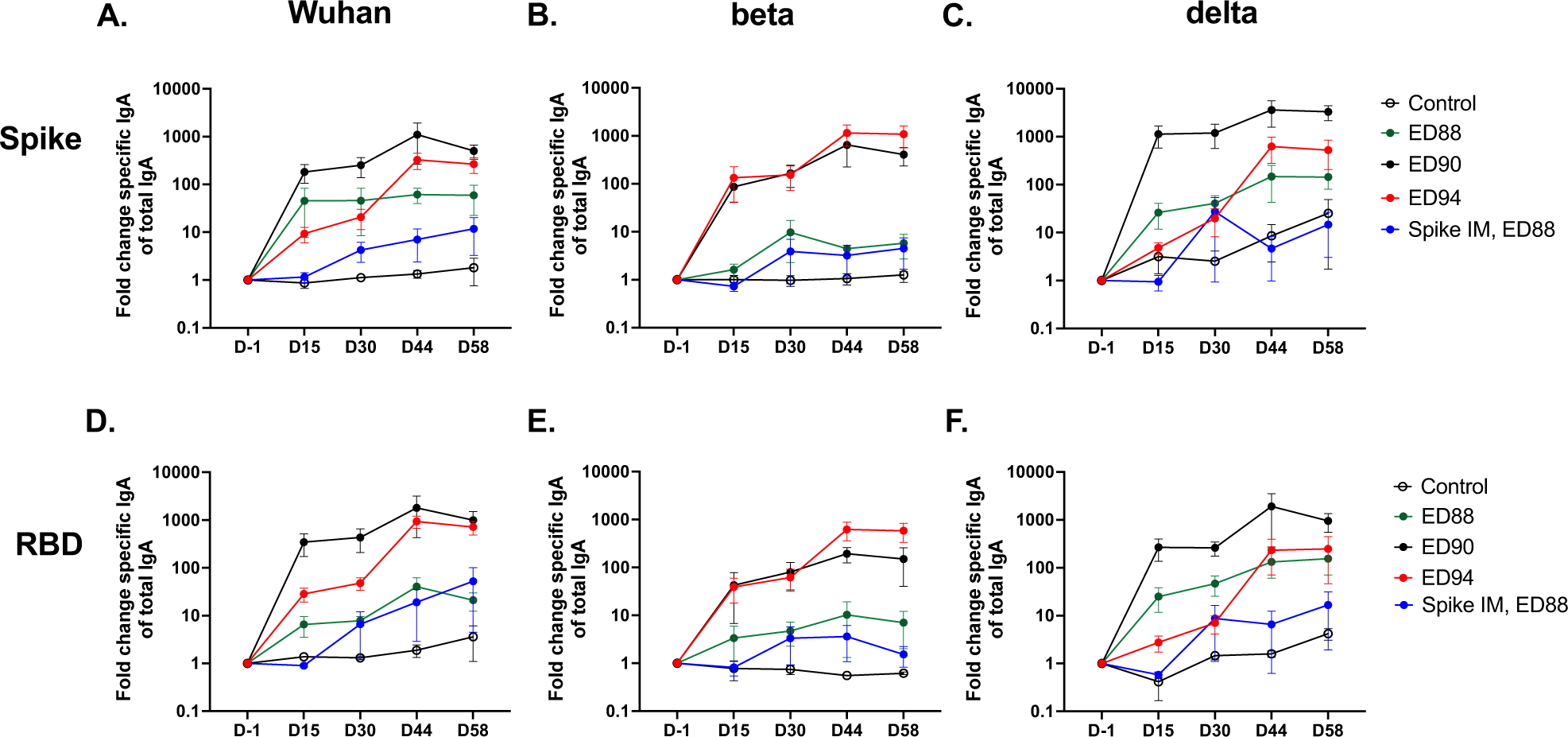
Immunization with ED90 greatly increases the production of nasal IgA to full length and RBD portions of S protein. IgA quantified on D-1, D15, D30, D44 and D58 to full length S-protein in nasal eluants **(A)** Wuhan **(B)** beta variant (B.1.351) and **(C)** delta variant (B.1.617.2). RBD specific IgA responses to **(D)** Wuhan **(E)** beta and **(F)** delta. Data normalized to total IgA in each sample timepoint and expressed as fold change from baseline at D-1, SEM (n = 4).

We investigated if the nasal SIgA specific to Wuhan and delta S protein had functional activity and could block the interaction with the host cell receptor ACE-2. The highest percent inhibition between ACE-2 and S protein was detected in the nasal samples from animals immunized with ED90 on D58 by surrogate virus neutralization test (sVNT) **(Figure 6)**. These results demonstrate that immunization in NHPs with vaccine candidate ED90 substantially elevates cross-reactive nasal SIgA that has blocking activity against Wuhan and delta S protein interactions with ACE-2.

**Figure 6:**
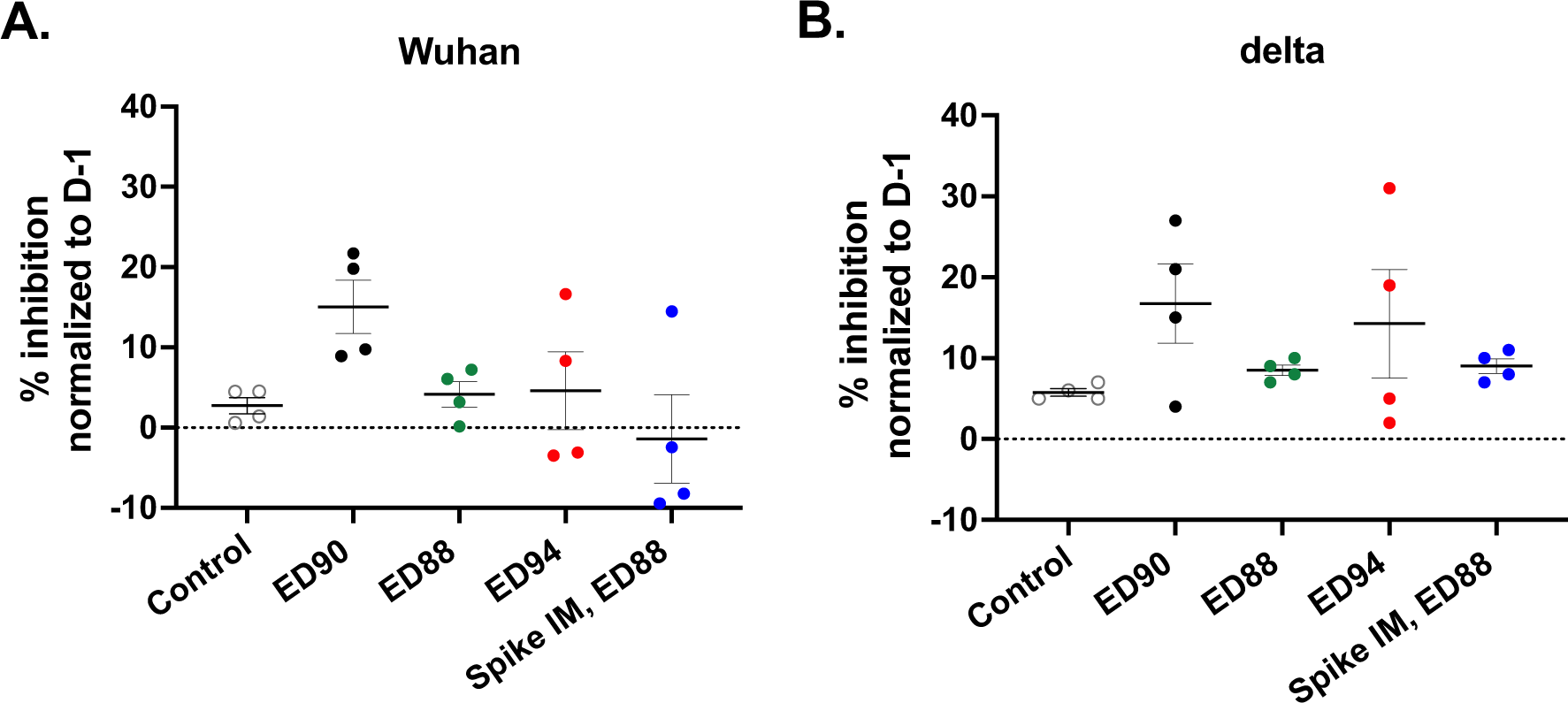
Immunization with ED90 increases neutralizing antibody in nasal samples to both Wuhan and delta by D58 post vaccination. percent inhibition of ACE2 binding in nasal samples to full length S protein **A)** Wuhan **(B)** delta variant (B.1.617.2) assessed by SVNT assay on D58. Data normalized to percent inhibition on D-1, SEM (n = 4).

## Conclusions

This study successfully identified clear differences in humoral immune responses in NHPs following mucosal administration of three candidate rAd5 vaccine constructs. The ED90 vaccine candidate appeared to be most potent construct at eliciting antibody responses to the original Wuhan SARS-CoV-2 strain and the most recent delta variant, without a substantial loss in activity to delta. Results in human trials with the mRNA vaccines suggest that the serum antibody responses to delta are reduced in comparison to Wuhan [3], therefore the data presented in this study suggests delivering rAd to a mucosal surface may enhance cross protective serum responses. Surprisingly the N protein in ED88 did not seem to induce substantial antibody responses to N (data not shown) and did not elicit as high of serum antibody titers to S protein and variants as ED90 did. When comparing the vaccine candidates that only express the S proteins, ED94 which express the beta VOC S protein, was better at inducing antibody responses to the homologous beta S protein, however, this vaccine candidate was inferior to the cross protective humoral responses generated by immunization with ED90 to the other S proteins tested. These results gave us confidence to go forward with ED90, rename it a clinical candidate designation VXA-CoV2-1.1-S, and start enrolling in a clinical trial (NTC05067933).

Current vaccines delivered under EUA protect against hospitalization and severe illness, but do not appear to be as effective at blocking infection or inhibiting transmission. In a study in Singapore by Chia, et al, vaccinated subjects were still capable of being infected by the delta variant at similar rates to unvaccinated subjects, with the vaccinated subjects able to clear the virus slightly faster [21]. In a Danish household study, it was determined that a full-vaccination regiment plus a booster was much more likely to prevent transmission of delta than omicron, with an odds ratio of 3.66 [22]. However, household transmission still occurred in 25-32% of the time even with a highly vaccinated population. These studies highlight that the prevention of infection and viral spread by current vaccine regimens is still a major hurdle towards ameliorating the large social, medical, and economic impact of the SARS-CoV-2 pandemic.

As each new variant appears, major outbreaks create substantial interference with normal life and economic activity. While omicron might not be as dangerous to vaccinated subjects as earlier variants, there are significant economic impacts due to infection. Infected workers miss work, strain testing centers, and impact emergency rooms. Major outbreaks could lead to closures of schools, gymnasiums, and public spaces. Although EUA guidelines for approval of a COVID-19 vaccine is the prevention of symptomatic disease, societal demands in the ongoing pandemic have shifted the goal posts to include prevention from infection and reduction of transmission. Interventions that induce antibodies in the nasal cavities may do a better job at preventing infection and reducing shedding/transmission for respiratory pathogens. For example in a Guinea pig influenza infection study, injected SIgA antibodies (that homed to the nasal mucosa) were able to prevent transmission from infected animals better than injected IgG antibodies[12]. Influenza viral shedding in humans has also been shown to be significantly lower when administered an oral vaccine that induced a mucosal IgA response, compared to individuals immunized with a commercial influenza vaccine, despite the commercial vaccine making 10 fold higher serum neutralizing antibody titers [23]. Furthermore, SARS-Cov-2 aerosol transmission to unvaccinated animals has been demonstrated to be significantly reduced in a vaccine breakthrough study when the ED90 vaccine candidate was given orally or intranasally [13].

In this currently study, the NHPs given rAd5 vaccines by the intranasal route induced substantial antibody responses in the nasal passages, which were highly cross-reactive against the S protein of several different variants. Additionally, we showed that both binding and neutralizing antibodies were found in the nasal passages of vaccinated monkeys, a first by any vaccine candidate. These results suggest that an effective mucosal response is possible and may lead to protection from infection, not just severe disease. We also determined the most potent approach for inducing a mucosal cross-reactive IgA response was found to be two doses of the ED90 vaccine candidate, which is now being evaluated in clinical trials.

## Supplemental figures

**Supplement figure 1:**
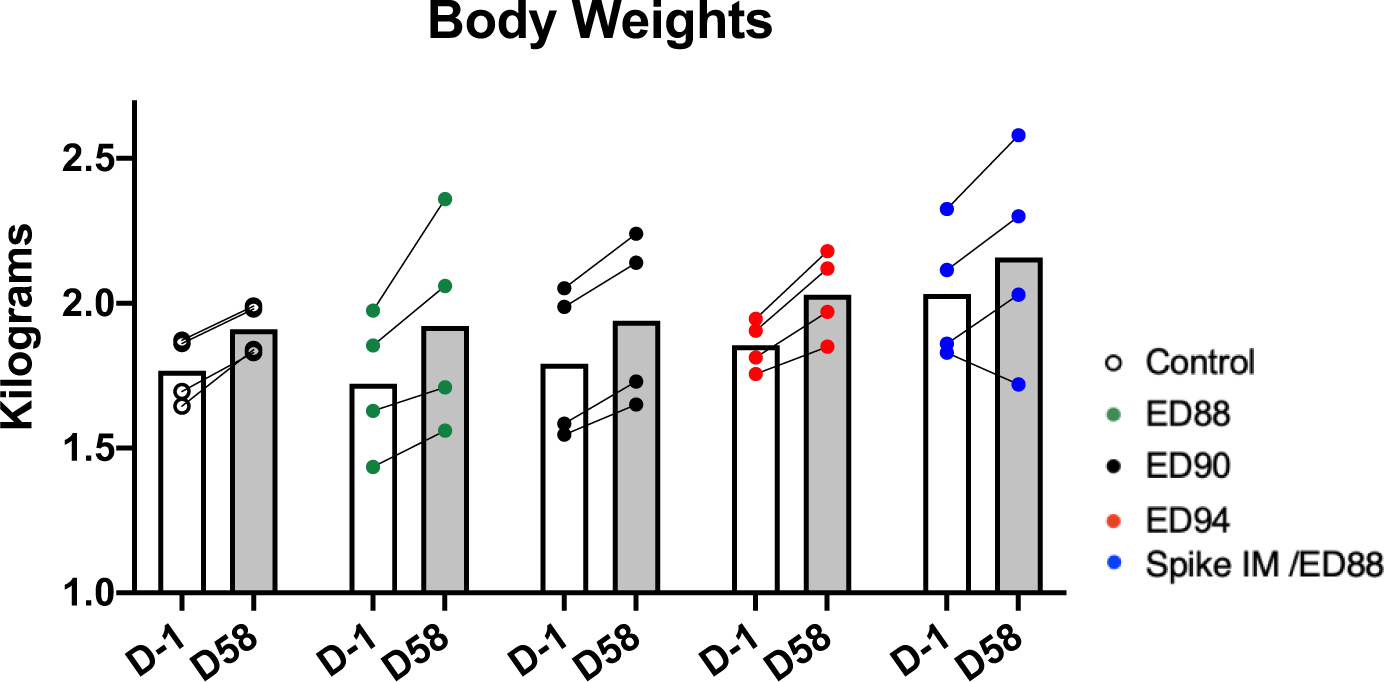
Animal body weights during immunogenicity study.

**Supplement figure 2:**
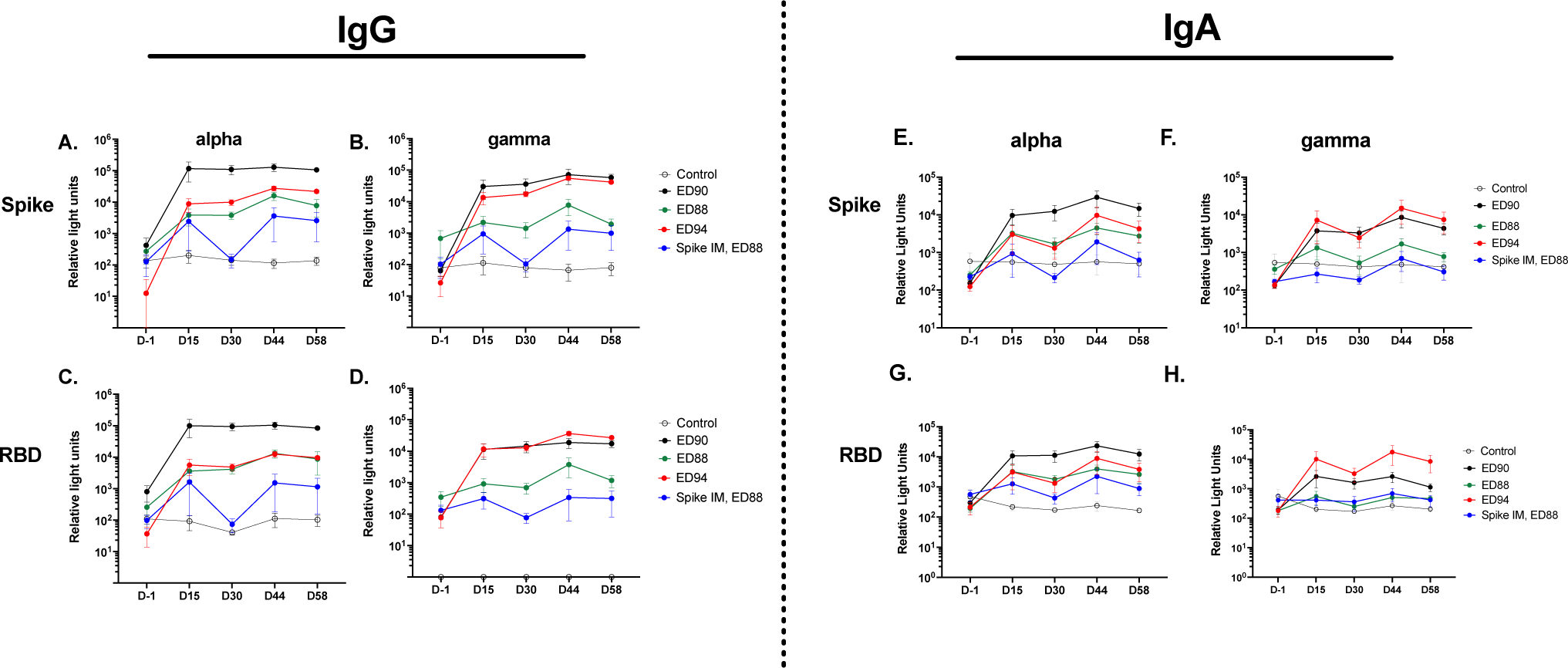
Serum IgG and IgA responses to full length S protein and RBD to alpha and gamma variants.

**Supplement figure 3:**
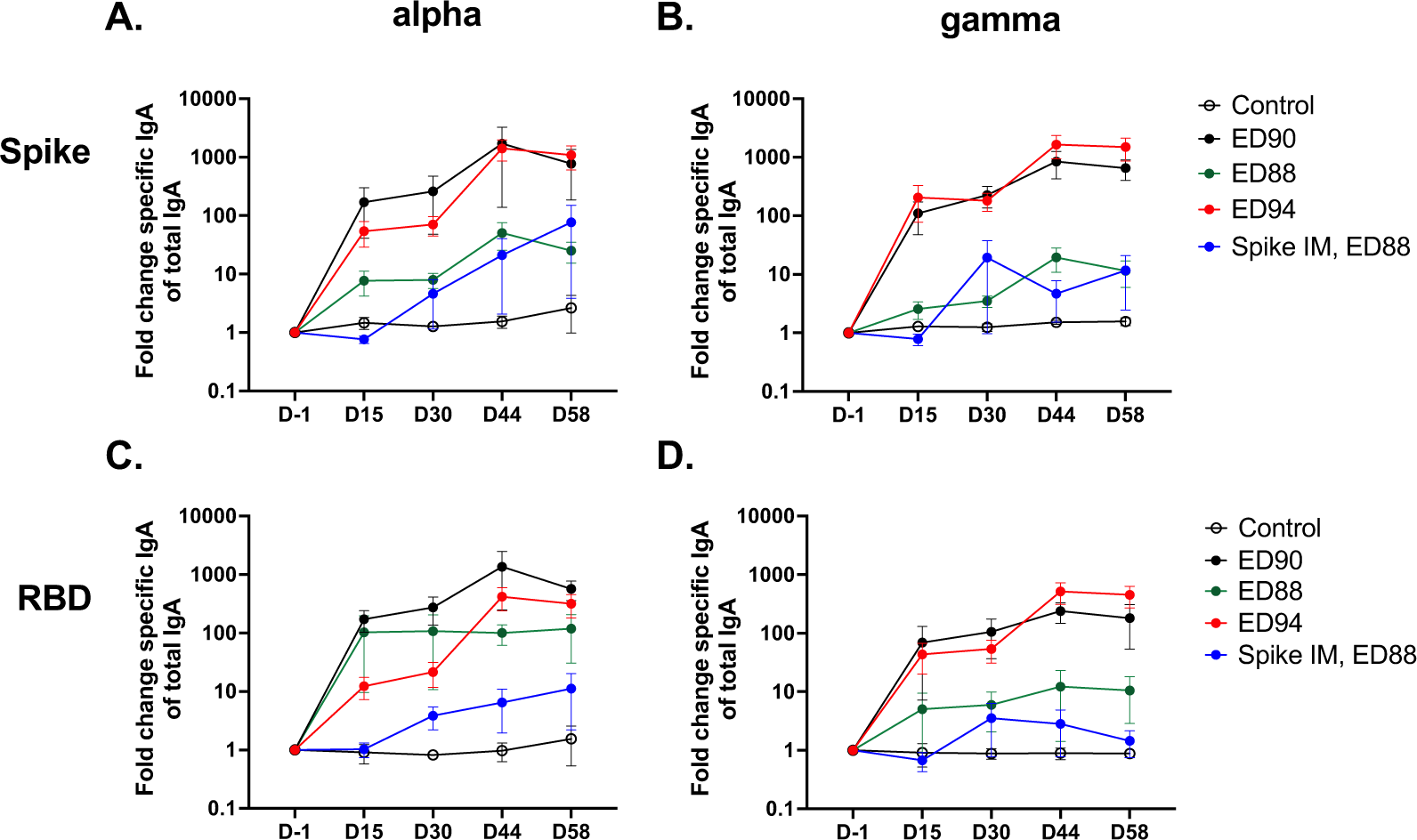
Nasal IgA responses to full length S protein and RBD to alpha and gamma variants.

## Methods

### Animals and study design

Twenty cynomolgus macaques were randomly assigned to five groups (*n =* 4 animals) comprising of two males and two females. Animals were immunized with rAd5 5×10^10^ infectious units (I.U.) on days 0 and 28 by intranasal administration using a mucosal atomization device (MAD) which delivered approximately 0.1 mL vaccine construct/nostril. Three groups were administered either ED88, ED90, or ED94 for prime and boost, and a fourth group was given an intramuscular injection of purified S protein, NR-52308 (BEI Resources) on day 0 followed by an intranasal boost of ED88 on D28. Serum and nasal swabs FLOQSwabs #5199CS01 (Copan Diagnostics, CA) were collected on day -1 from each animal prior to immunization and on day 15, 29, 44 and 58 post vaccination (+/-24 hrs). Animals were monitored daily for any abnormal clinical observations and body weights recorded weekly.

### Adenoviral vaccine constructs

Adenovirus Type 5 (rAd5) vaccine constructs were created based on the published DNA sequence of SARS-CoV-2 publicly available as Genbank Accession No. MN908947.3. The published amino acid sequences of the SARS-CoV-2 S or N were used to create recombinant plasmids containing transgenes cloned into the E1 region of rAd5 [24], using the same vector backbone used in prior clinical trials for oral rAd tablets [14, 25]. All vaccines were grown in the Expi293F suspension cell-line (Thermo Fisher Scientific) and purified by CsCl density centrifugation.

### Serum IgG and IgA antibody responses against SARS-CoV-2 variants

Serum IgG and IgA to responses to full length S protein and RBD was quantified against Wuhan and alpha, beta and gamma variants, using V-PLEX SARS-CoV-2 panel 7 (Meso Scale Diagnostics, MD). A 2-sector U-plex (Meso Scale Diagnostics, MD) assay was developed to measure delta variant specific serum IgG and IgA. 66nM of biotinylated full length trimerized delta spike (L452R, T478K) and RBD (Acro Biosystems, DE) was conjugated to 2 different linkers and multiplexed on a 2-sector U-plex plate according to manufacture protocols. For both V-plex or U-plex assays, serum was diluted 1:5000 in 1% ECL Blocking Agent (Cytiva, United Kingdom) in 1X PBS, 0.05% Tween 20 and assayed in duplicate and incubated room temperature for two hours shaking at 700rpm. An 8-point reference serum using reference standard 1 (Meso Scale Diagnostics, MD) was serially diluted 4-fold and included in all assays starting at 1:10. Plates were washed with 1X PBS, 0.05% Tween 20 and incubated with 1:200 dilution of 200X Sulfo-Tag anti-IgG or IgA (Meso Scale Diagnostics, MD) for 1 hour shaking at 700rpm. Plates were developed using MSD Gold Read Buffer and data acquired using the MSD Sector Imager 120 instrument.

### Nasal IgA responses to SARS-CoV-2 variants

Nasal secretions were collected from right and left nostrils using two FLOQSwabs #519CS01 (Copan Diagnostics, CA), and were immediately frozen and stored at -80°C. To elute the antibodies the swabs were thawed at 20-22°C for 15 minutes and transferred into Eppendorf tubes containing 300 uL elution buffer (0.05% Tween, 1% BSA, 1X PBS). The swab tips were cut and vortexed inside the Eppendorf tubes for 30 seconds and the resulting liquid was transferred into pre-charged elution columns (Costar Spin-X, Corning). The elution columns were spun to remove debris at 16,000 g at 4°C for 20 mins and the left and right nasal eluants were then pooled, aliquoted and frozen at -80°C prior to use. To measure IgA in the nasal samples, eluants were diluted 1:4 and 1:16 in 1% ECL Blocking Agent (Cytiva, United Kingdom) and run on a V-PLEX SARS-CoV-2 (Panel 7) or 2-sector U-plex assay (Meso Scale Diagnostics, MD) described above. Antigen-specific nasal IgA was normalized to the total amount of IgA in the corresponding sample. Fold change was calculated by dividing the normalized antigen-specific nasal IgA values in each timepoint by the D -1 values.

### Serum PRNT neutralizing assay

PRNT assays were conducted at Bioqual as described [20]. Vero E6 cells (ATCC, cat# CRL-1586) were plated in 24-well plates at 175,000 cells/well in DMEM + 10% FBS + Gentamicin, until cells reached 80 -100% confluency the following day. Serum samples were heat inactivated at 56°C for 30 minutes and serially 3-fold diluted with a starting dilution of 1:10 in a dilution plate until use. A 30 pfu/well concentration of the WA/2020 (other 2 strains) virus was prepared and 300 μL was added to all samples and positive control wells. This doubled the sample dilution factor (1st well begins at 1:20). The plate was then covered with a plate sealer and incubated at 37°C, 5.0% CO_2_ for 1 hour. The media from the 24-well plate was removed and 250 μL of serum and virus mixed titrated samples was added in duplicate to the Vero E6 cells. The 24-well plates were incubated at 37°C, 5.0% CO_2_ for 1 hour for virus infection. 0.5% methylcellulose media was heated in a 37°C water bath and 1 ml was added to each well and the plates then incubated at 37°C, 5% CO_2_ for 3 days. The methylcellulose medium was removed, and the plates were washed once with 1 mL PBS and fixed with 400μL ice cold methanol per well at -20°C for 30 minutes. After fixation, monolayers were stained with 250 μL per well of 0.2% crystal violet (20% MeOH, 80% dH_2_O) for 30 minutes at room temp and then washed once with dH_2_O and left to dry for at least 15 min. The plaques in each well were recorded and the IC_50_ and IC_90_ titers were calculated based on the average number of plaques detected in the virus control wells. A control (rabbit) reference serum with established titer (5,400 IC_50_) was included in each assay set-up to serve as an internal positive control.

### Nasal secretion SVNT

Neutralizing antibodies in nasal secretions to SARS-CoV-2 spike and RBD (Wuhan) were measured by MSD V-PLEX SARS-CoV-2 Panel 2 (Meso Scale Diagnostics, MD) kits according to the manufacturer’s protocol. Blocking antibodies to delta spike and RBD were detected with MSD U-Plex Development Pack 2-spot kit. Nasal samples were diluted of 1:4 and 1:16 in MSD diluent 100 and incubated for 1 hour at room temperature shaking at 700rpm.

Human ACE2 protein conjugated with SULFO-TAG label was added to wells and incubated for 1 hour at room temperature with shaking at 700rpm. Plates were developed using MSD Gold Read Buffer and data acquired using the MSD Sector Imager 120 instrument. Results were reported as percent inhibition (1-(average sample ECL signal / average ECL signal of blank well) x 100).

